# Efficient Permutation-based Genome-wide Association Studies for Normal and Skewed Phenotypic Distributions

**DOI:** 10.1101/2022.04.05.487185

**Authors:** Maura John, Markus J Ankenbrand, Carolin Artmann, Jan A Freudenthal, Arthur Korte, Dominik G Grimm

## Abstract

**Motivation:** Genome-wide Association Studies (GWAS) are an integral tool for studying the architecture of complex genotype and phenotype relationships. Linear Mixed Models (LMMs) are commonly used to detect associations between genetic markers and the trait of interest, while at the same time allowing to account for population structure and cryptic relatedness. Assumptions of LMMs include a normal distribution of the residuals and that the genetic markers are independent and identically distributed - both assumptions are often violated in real data. Permutation-based methods can help to overcome some of these limitations and provide more realistic thresholds for the discovery of true associations. Still, in practice they are rarely implemented due to its high computational complexity.

**Results:** We propose `permGWAS`, an efficient linear mixed model reformulation based on 4D-tensors that can provide permutation-based significance thresholds. We show that our method outperforms current state-of-the-art LMMs with respect to runtime and that a permutation-based threshold has a lower false discovery rate for skewed phenotypes compared to the commonly used Bonferroni threshold. Furthermore, using `permGWAS` we re-analysed more than 500 *Arabidopsis thaliana* phenotypes with 100 permutations each in less than eight days on a single GPU. Our re-analyses suggest that applying a permutation-based threshold can improve and refine the interpretation of GWAS results.

**Availability:** `permGWAS` is open-source and publicly available on GitHub for download: https://github.com/grimmlab/permGWAS.

## 1 Introduction

Genome-wide Association Studies (GWAS) are an integral tool to detect associations between genetic markers within a population of individuals and a complex trait or disease that is measured in the same population (Atwell *et al*., 2010; The 1001 Genomes Consortium, 2016; Todesco *et al*., 2020). State-of-the-art methods can easily analyze millions of markers in populations of thousands of individuals (Kang *et al*., 2008, 2010; Lippert *et al*., 2011; Korte *et al*., 2012; Loh *et al*., 2015). Here, the critical step is to define a threshold to distinguish true and spurious associations. Classically, one controls the family-wise error rate (FWER), that is the probability of making at least one type-1 error (or false positive), using the commonly used Bonferroni correction (Bonferroni, 1936). However, due to the large number of tests the Bonferroni correction is in practice often too conservative (Westfall and Young, 1993; Llinares-López *et al*., 2015; Gumpinger *et al*., 2021), as it assumes that all tested markers are independent, which is clearly not the case for high-density genomic data that are nowadays routinely generated. Here, many markers are correlated with each other and the actual number of independent tests performed is lower than the number of markers analyzed. Therefore, many studies propose a significance threshold that is based on the false-discovery rate (FDR) (Storey and Tibshirani, 2003). On the other hand, naïve thresholds, such as Bonferroni or FDR, cannot account for model miss-specifications that easily arise in biological data, which are often non-normally distributed. Variance-stabilizing transformations have been proposed to account for phenotypic variability (Sun *et al*., 2013), but are not non-controversial (Shen and Rönnegård, 2013) and might complicate comparability across different phenotypes.

Permutation-based thresholds could provide an alternative approach to overcome some of these limitations (Che *et al*., 2014). Here, the main limitations are the computational burden to run permutations routinely, as current implementations are still too slow and inefficient (such as our deprecated method `GWAS-Flow` (Freudenthal *et al*., 2019)), or focus only on linear regression without the possibility to correct for confounding factors on specialised FPGA (Field Programmable Gate Arrays) hardware (Swiel *et al*., 2022). We propose `permGWAS`, an efficient permutation-based linear mixed model to compute adjusted significance thresholds that are able to account for correlated markers and skewed phenotypic distributions without the need to arbitrarily transform phenotypes. To account for multiple hypothesis, correlated markers and skewed phenotypes, we compute permutation-based significance thresholds based on the *maxT* methods proposed by Westfall & Young (Westfall and Young, 1993). To enable efficient computation of different permutation-based tests, we provide a scale-able batch-wise reformulation of a permutation-based linear mixed model using 4 dimensional tensors. We propose to implement permutation-based thresholds as the default choice for GWAS and provide both simulations and re-analysis of more than 500 *Arabidopsis thaliana* phenotypes to underpin its benefits.

## 2 Methods

We will first provide the necessary background of linear mixed models for genome-wide association studies, the multiple hypothesis testing problem and how to empirically estimate the family-wise error rate using the Westfall-Young permutation testing procedure (Westfall and Young, 1993). Finally, we will present our approach on how to efficiently compute associations with linear mixed models (LMMs) using a permutation-based significance threshold. An overview of all mathematical symbols and notations can be found in Suppl. Tab. 1.

### 2.1 Linear Mixed Model

Let *n* be the number of samples and *m* the number of genetic markers. Then for each genetic marker we consider a LMM of the form

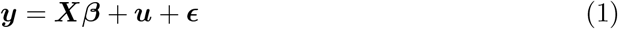

where 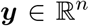 is a vector of observed phenotypic values and 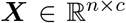 is a matrix of fixed effects containing columns for the mean, covariates and the genetic marker. Fixed effects are denoted by 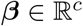 and random effects 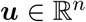 follow a Gaussian distribution with zero mean and a (genetic) variance of 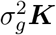, where 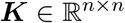 denotes the kinship matrix and 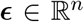 is a vector of residual effects with 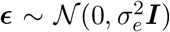. As described in (Kang *et al*., 2008, 2010; Lippert *et al*., 2011), we estimate the variance components 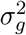 and 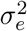 by maximizing the following likelihood function

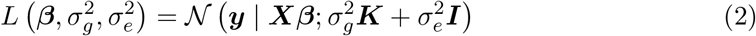

for both, the null model which includes no genetic markers and afterwards the alternative model which includes the marker of interest. Finally, a F-test is used to test the null hypothesis that the marker has no effect against the alternative hypothesis that it has an effect on the phenotypic value. We can reject the null hypothesis and call a statistical test significant, if the p-value of the F-test is below a predefined significance threshold *α* (e.g. 5%).

### 2.2 Multiple Hypothesis Testing

Since we have to test thousands to millions of markers simultaneously, we have to take these multiple tests into account, otherwise we would obtain thousands of false positive associations deemed to be significant.

#### 2.2.1 Family-Wise Error Rate

The family-wise error rate (FWER) is the probability of making at least one type-1 error (or false positive). One has to find an appropriate corrected significance threshold *δ* for each hypothesis, such that the FWER(*δ*) ≤ *α*. To determine the optimal threshold *δ*\* one has to solve the following optimization problem:

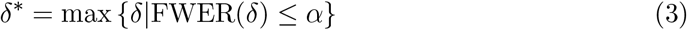

Evaluating this optimization problem in closed form is not possible in general. For this purpose, the widely used Bonferroni approximation (Bonferroni, 1936) can be used to control the FWER. To estimate the adjusted significance threshold 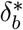 after Bonferroni, one simply divides the number of simultaneous tests by the target significance level *α*, i.e. 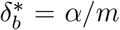. However, due to the large number of tests the Bonferroni correction is in practice often too conservative, i.e. 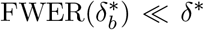, as shown in (Llinares-López *et al*., 2015; Gumpinger *et al*., 2021). In addition, when performing GWAS one typically makes the assumption that the residuals are normally distributed and that the genetic markers are independent and identically distributed. However, these assumptions are often violated in practice, which leads to the fact that the Bonferroni threshold is either overly conservative for normally distributed phenotypes (leading to many false negatives) or not stringent enough for phenotypes with skewed distributions (leading to many false positives).

#### 2.2.2 Westfall-Young Permutations

Permutation-based methods can help to overcome some of these problems, by empirically estimating the FWER(*δ*). One could either approximate the null distribution by using permutations to then compute adjusted p-values or to use the unadjusted p-values and provide a permutation-based significance threshold based on the *maxT* permutation-method proposed by Westfall and Young (Westfall and Young, 1993). With this adjusted threshold we can account for non Gaussian distributed phenotypes, correlated markers due to linkage disequilibrium (LD) and the large number of tests. In the following we will describe how to compute both, adjusted p-values and adjusted significance thresholds. To compute adjusted p-values, we first permute the phenotype *q* times and calculate the test statistics ^(*k*)^*t_j_* for the *k*^th^ permutation, with *k* ∈ {1,…, *q*} and *j*^th^ marker, with *j* ∈ {1,…, *m*}. After randomizing, any correlation left between the genotypic and phenotypic values will be of non-genetic origin, but the distribution of the phenotypic values stays the same. To compute the permutation-based p-values, let *T_j_* denote the random variable corresponding to the observed test statistic of the *j*^th^ marker. We test the hypothesis *H*_0_ that *T_j_* follows the permutation distribution empirically given by ^(*k*)^*t_j_* for all *k* and all *j*. Then we compute the adjusted permutation-based p-value as:

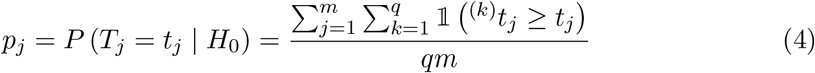

Where 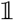 takes the value 1 if the argument is true and 0 otherwise. The FWER can be controlled in this multiple hypothesis testing setting using Bonferroni (Bonferroni, 1936).

For the adjusted significance threshold we follow a permutation testing procedure proposed by Westfall and Young (Westfall and Young, 1993). For each permutation we take the maximal test statistic over all markers, ^(*k*)^*t_max_* = max_*j*∈{1,…, *m*}_ ^(*k*)^*t_j_* and compute the corresponding minimal p-value ^(*k*)^*p_min_*. Let again *T_j_* denote the random variable corresponding to the observed test statistic of the *j*^th^ marker. We now test the hypothesis *H*_0_ that *T_j_* follows the permutation distribution empirically given by ^(*k*)^*t_max_* for all *k*. Then the adjusted p-value is given by 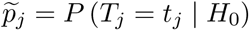 and

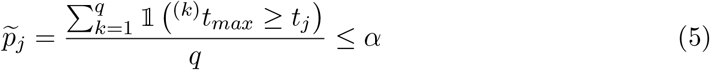

is equivalent to *t_j_* being larger than the 100(1 – *α*)^th^ percentile of the ^(*k*)^*t_max_*. Hence, the *α*^th^ percentile of the minimal p-values ^(*k*)^*p_min_* leaves us with an adjusted threshold that controls the FWER.

### 2.3 permGWAS Architecture

These permutation-based strategies are computationally highly demanding, which makes them often inapplicable in practise. Further, current state-of-the-art GWAS implementations sequentially compute univariate test-statistics for one marker and a given phenotype (Kang *et al*., 2008, 2010; Lippert *et al*., 2011; Grimm *et al*., 2017). We propose `permGWAS`, which is able to simultaneously compute univariate test statistics of several SNPs batch-wise on modern multi-CPU and GPU environments, while at the same time controlling the FWER using Westfall-Young permutation testing. First, we will introduce the mathematical framework for batch-wise linear mixed models without permutations, followed by an efficient formulation for permutation-based linear mixed models.

#### 2.3.1 Batch-Wise Linear Mixed Models

Denote by *n* the number of samples, c the number of fixed effects (i.e. the SNP of interest and all covariates) and *b* the batch size. Let 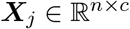 denote the matrix of fixed effects, including a column of ones for the intercept, the covariates and the *j*^th^ SNP 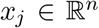 (Fig. 1A). Let 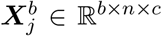 be the 3D tensor containing the matrices ***X**_j_* to ***X***_*j*+*b*−1_ and let 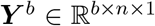 denote the 3D tensor containing b copies of the phenotype vector 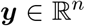 (Fig. 1B). Further, let 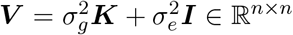 denote the variance-covariance matrix. For computational efficiency, instead of using generalized least squares, we first compute the Cholesky decomposition ***V*** = ***CC**^T^* and linearly transform 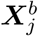 and ***Y**^b^*, before computing the coefficients using ordinary least squares. Let 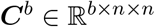 denote the 3D tensor containing *b* copies of ***C***. Then the linearly transformed data is given by

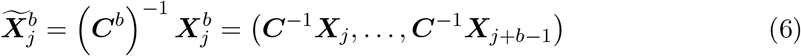

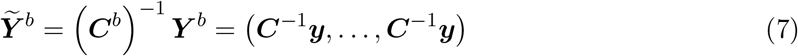

**Figure 1:**
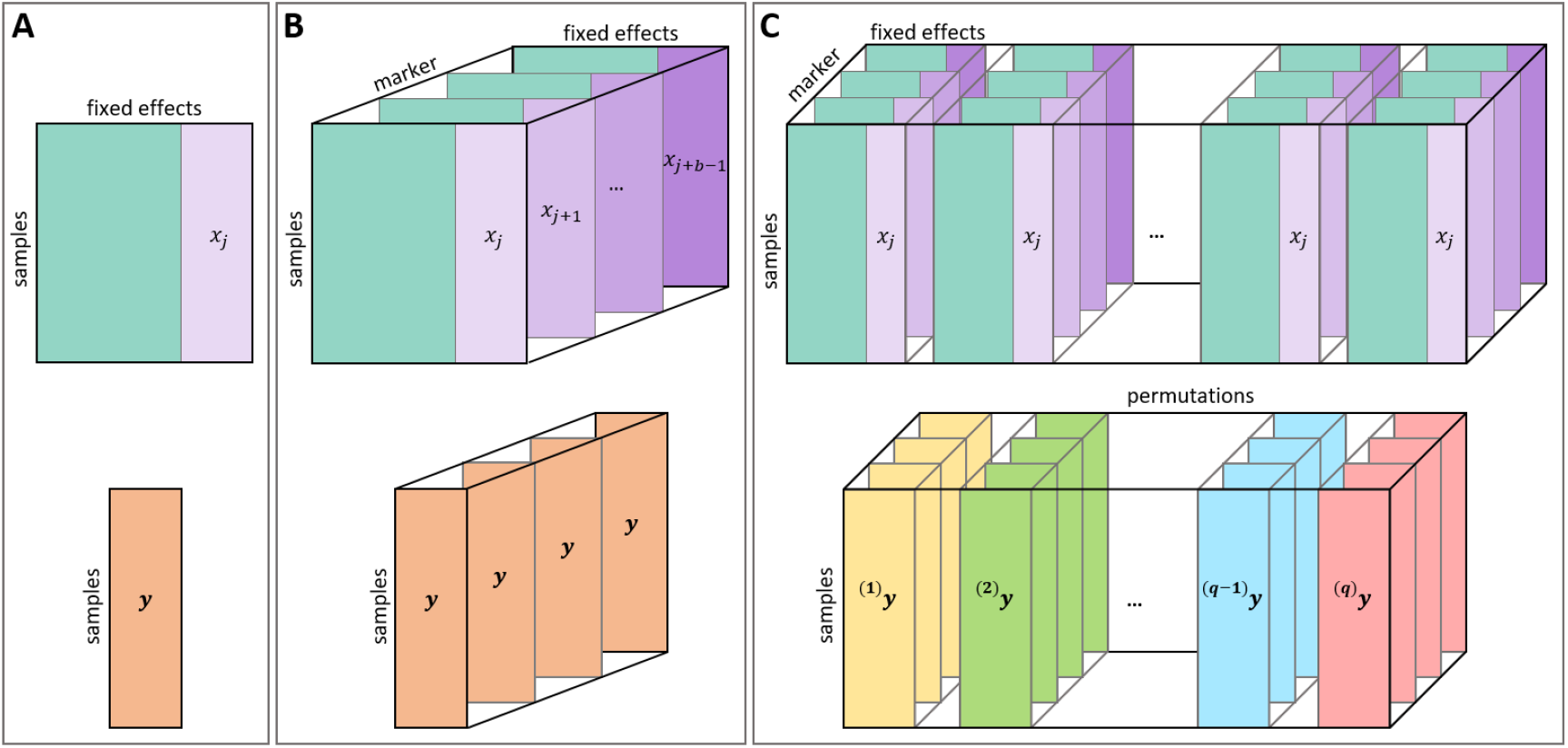
Schematic illustration of matrices and tensors of the `permGWAS` architecture. (A) Commonly used matrix representation when computing sequential univariate tests, where 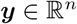 is the phenotypic vector for *n* samples and 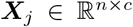 denotes the matrix of fixed effects, including a column of ones for the intercept, the covariates and the *j*^th^ SNP 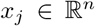. (B) 3D-tensor representation of a LMM to compute univariate tests batch-wise. The phenotype is represented as a 3D tensor containing *b* copies of the phenotype vector 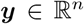 and 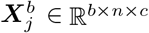 is a 3D tensor containing the matrices ***X**_j_* to ***X***_*j*+*b*-1_. (C) 4D-tensor representation of a permutation-based batch-wise LMM. The phenotype is represented as a 4D tensor containing for each permutation ^(*k*)^***y*** the 3D tensor ^(*k*)^***Y***^*b*^ for all *q* permutations and 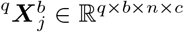 a 4D tensor containing *q* copies of 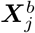

Now we can compute the coefficients 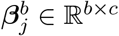 and the residual sums of squares 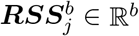 for those *b* SNPs starting at the *j*^th^ SNP as follows:

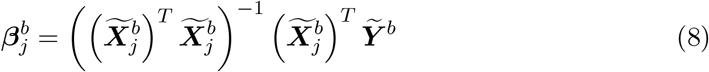

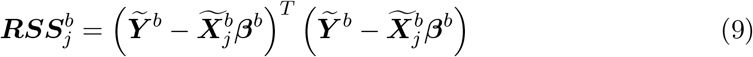

Where 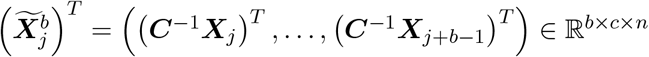. Finally, we can compute the test statistics 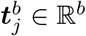 for all *b*-SNPs:

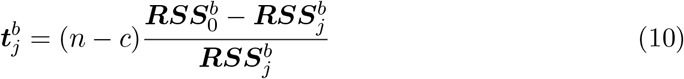

where 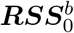 contains *b* copies of the residual sum of squares of the null model. Once we have computed the test statistics for all SNPs we can sequentially calculate all p-values.

#### 2.3.2 Efficient Permutation-Based Linear Mixed Models

When performing GWAS with permutations let additionally q denote the number of permutations. Then for each permutation ^(*k*)^*y* of *y* with *k* ∈ {1,…, *q*} we get a new 3D tensor 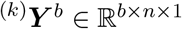. Let 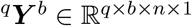 be the 4D tensor containing ^(*k*)^***Y**^b^* for all *k* and let 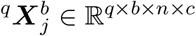 be the 4D tensor containing *q* copies of 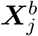 (Fig. 1C). Now for each permutation ^(*k*)^*y* of *y* we estimate associated variance components 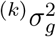 and 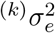 and obtain a new variance-covariance matrix 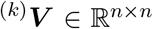. We compute the Cholesky decomposition ^(*k*)^***V*** = ^(*k*)^***C*** ^(*k*)^***C**^T^* for each *k* and again linearly transform the data. Let 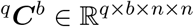 denote the 4D tensor containing the 3D tensors 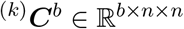 for all *k*. Then we can transform the data via

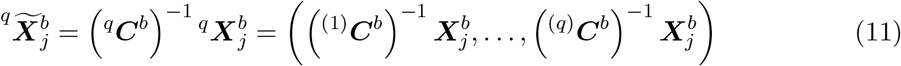

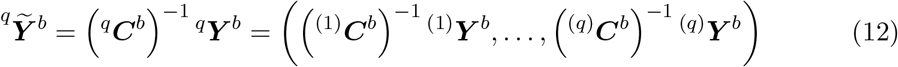

Now similar to above we compute the coefficients 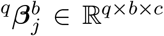, the residual sums of squares 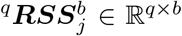 and the test statistics 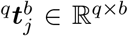 for all *q* permutations and *b* SNPs at once:

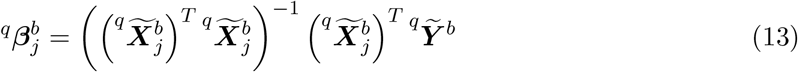

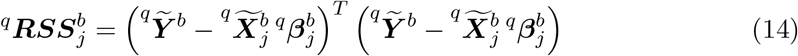

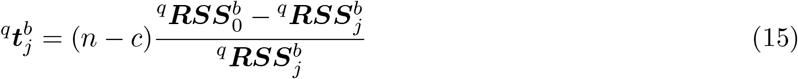

Where

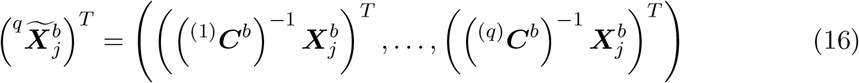

is a 4D tensor in 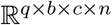 and 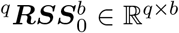 contains *b* copies of the RSS of the null model for each permutation.

### 2.4 Implementation

The `permGWAS` framework is implemented in Python3 using commonly used libraries for scientific computing, such as `numpy` (Harris *et al*., 2020), scipy (Virtanen *et al*., 2020), pandas (McKinney *et al*., 2011) and `PyTorch` (Paszke *et al*., 2019) to support efficient tensor arithmetic as well as multi-core and GPU support. In addition, specialized packages for estimating the variance components (`limix` (Lippert *et al*., 2014)) and file IO (`h5py, pandas-plink`) are used. `permGWAS` can be used as a standalone command line tool or directly within Python. To ensure a smooth experience on different environments and machines we provide a standardized Docker environment. Our Framework supports several common genotype and phenotype file formats, including HDF5, CSV and PLINK (Purcell *et al*., 2007). Further, `permGWAS` supports to filter for minor allele frequency (MAF) and also to include one or more covariates to account for certain fixed effects. By default `permGWAS` computes as a kinship matrix the realized relationship kernel (Hayes *et al*., 2009), however it is also possible to provide any other type of genetic similarity matrix. In order to run the tool on different machines, the batch size for the simultaneous computation of univariate tests as well as the batch size for permutation-based tests can be adjusted. To reduce the memory footprint, it is also possible to load genotypic data continuously in chunks from a HDF5 file, in case a pre-computed kinship matrix is provided. All code is open-source and publicly available on GitHub, including more details and information on how to run the tool: https://github.com/grimmlab/permGWAS.

### 2.5 Data & Simulations

We evaluate the performance and runtime of `permGWAS` on simulated data as well as on publicly available genotype and phenotype data from the model plant Arabidopsis thaliana.

#### 2.5.1 Arabidopsis thaliana Data

As genotypic data a fully imputed SNP-Matrix of 2029 accessions and approximately 3M segregating markers is used (Arouisse *et al*., 2020). Phenotypic data for 516 different traits were downloaded from the central and manually curated AraPheno database (Seren *et al*., 2016; Togninalli *et al*., 2020).

#### 2.5.2 Simulations

Artificial phenotypes were simulated for 200 random *Arabidopsis thaliana* accessions using the fully imputed SNP matrix from above. For each synthetic phenotype 1 001 SNPs with a minor allele frequency of 5% or higher were randomly sampled, where 1 SNP was considered causative and the other 1 000 were used to simulate the polygenic background. Here, each background SNP contributed a small random amount, drawn from a normal distribution with *μ* = 0 and *σ* = 0.1 to the phenotypic value. Random noise drawn from a gamma or normal distribution was added, such that the noise accounts for 70% of the total phenotypic variance. Finally, a fixed effect for the causative SNP was added to explain roughly 20% of the total genetic variance. In this manner, six different sets containing 50 phenotypes each, were simulated. The sets differed by the distribution of the noise, where one set had normally distributed noise and the other five sets used gamma distributed noise with shape parameters of 0.1, 1, 2, 3 and 4. For evaluation, `permGWAS` was applied with 100 permutations on each of the 300 simulated phenotypes. Each phenotype was classified as true positive (TP) if any SNP in a 50kbp window around the causative marker was significant. Additionally, each phenotype was classified as false positive (FP) if any SNP outside the 50kbp window around the causative marker was significant. This way a phenotype can be true positive and false positive at the same time. We define the phenotype-wise false discovery rate (FDR) as 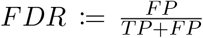. These values were calculated separately for the p-value thresholds based on both, the Bonferroni and permutations-based thresholds.

## 3 Results & Discussion

In the following we evaluate `permGWAS` with respect to runtime and statistical power using simulated data, as well as on more than 500 public available phenotypes from the model species *Arabidopsis thaliana*.

### 3.1 Results on Synthetic Data

#### 3.1.1 Runtime Comparisons

We analyzed the runtime of `permGWAS` with respect to (1) the number of markers, (2) the number of samples and (3) the number of permutations. For all runtime experiments we used data from a flowering time related phenotype in *Arabidopsis thaliana*, FT10 (flowering time at 10 degrees; DOI:10.21958/phenotype:261) (The 1001 Genomes Consortium, 2016), and down- and up-sampled the phenotype and corresponding SNP matrix to generate synthetic data with varying number of samples. We compared the runtime of `permGWAS` with two state-of-the-art and commonly used LMMs, `EMMAX` (Kang *et al*., 2010) and `FaST-LMM` (Lippert *et al*., 2011). For both, we used the binary C/C++ implementations. All runtime experiments were conducted on the same machine running Ubuntu 20.04.3 LTS with a total of 52 CPUs, 756GB of memory and 4 NVIDIA GeForce RTX 3090 GPUs, each with 24GB of memory. For our experiments we restricted the number of CPUs to 1 and 8 cores and a single GPU using dedicated Docker containers. We took the mean of the runtime over three runs for each experiment.

First, we compared the runtime on environments with a single CPU and GPU. For this purpose, we fixed the number of samples to 1000 and varied the markers between 10^4^ and 5 · 10^6^ to evaluate the effect of an increasing number of SNPs. As summarized in Fig. 2A all models show a linear dependency with respect to the number of SNPs. `permGWAS` (the GPU and CPU version) outperform both, the binary implementation of `EMMAX` and `FaST-LMM`. Our dockerized Python implementation of `permGWAS` is almost one order of magnitude faster than the C/C++ implementation of `FaST-LMM` (for 1000 samples and 5 · 10^6^ markers 0.33h and 2.8h, respectively). This can be mainly explained due to the batch-wise computation of several univariate statistical simultaneously.

**Figure 2:**
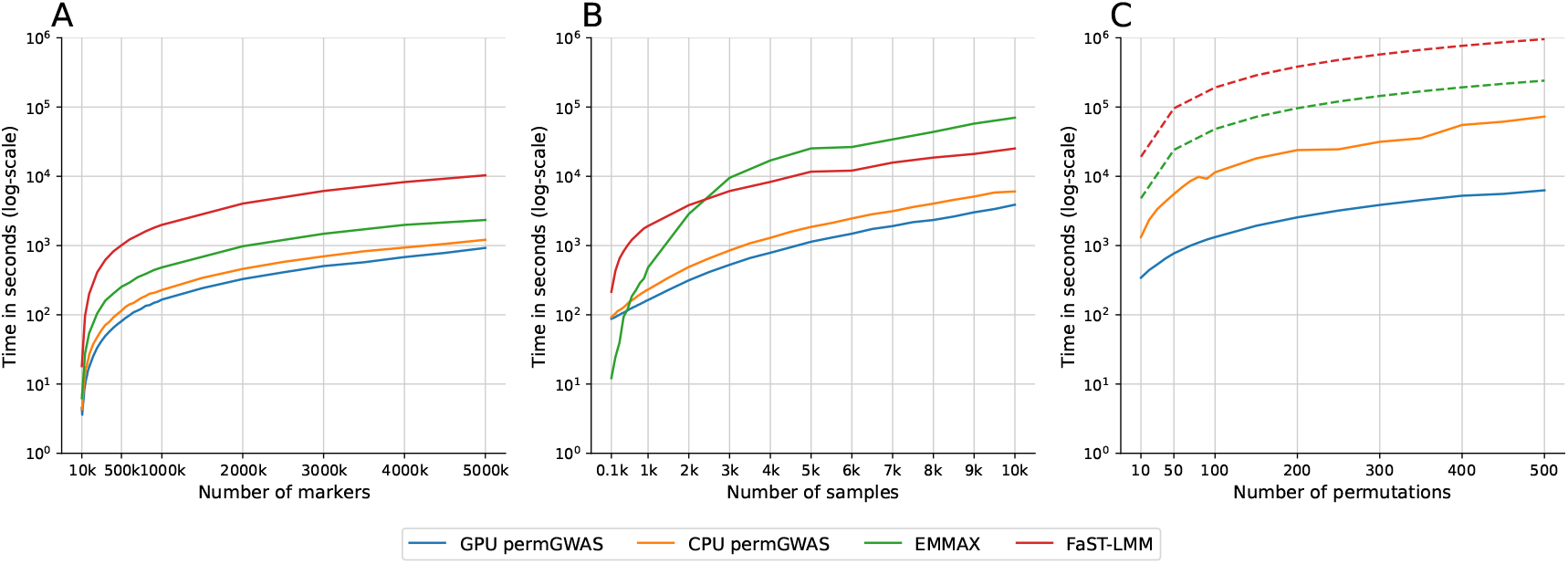
Runtime comparison of permGWAS vs. EMMAX and FaST-LMM. Note all axes are log-scaled. (A) Computational time as function of number of SNPs with fixed number of 1 000 samples. (B) Computational time as function of number of samples with 10^6^ marker each. (C) Computational time as function of number of permutations with 1 000 samples and 10^6^ marker each. Dashed lines for `EMMAX` and `FaST-LMM` are estimated based on the computational time for 1 000 samples and 10^6^ markers times the number of permutations.

Next, to estimate the effect of the number of samples on the runtime we fixed the number of SNPs to 10^6^ and varied the number of samples between 100 and 10^4^. In Fig. 2B we can observe that `EMMAX` outperforms all other comparison partner for sample sizes smaller than 500. However, the runtime increases quickly for larger samples sizes. Again `permGWAS` outperforms both comparisons partners by at least one order of magnitude. Here the runtime of the GPU version of `permGWAS` for 10^4^ samples and 10^6^ markers was approximately 1.7h, while for `FaST-LMM` and `EMMAX` the runtime was more than 7h and 19h, respectively. Finally, to compare the runtime of the permutation-based approach, we fixed the number of samples to 1000 and the number of SNPs to 10^6^ and conducted between 10 and 500 permutations with `permGWAS` using a single GPU architecture vs. a single CPU. Since `EMMAX` and `FaST-LMM` only perform one univariate test at a time and are not designed for permutation-based tests, we took the runtime for 1 000 samples and 10^6^ markers from the previous experiment and estimated the runtime for permutations by multiplying with the number of permutations. This is just an estimate of the minimal runtime, since no data pre-processing and post-processing steps are included (e.g. preparing permuted phenotypes, merging result files to estimate adjusted p-values/thresholds). The advantage of the GPU architecture becomes most obvious when using permutations, as illustrated in Fig. 2C. The GPU version of `permGWAS` is at least an order of magnitude faster than the CPU version of `permGWAS`. More importantly, `permGWAS` (GPU) is more than one order of magnitude faster than `EMMAX` and more than two orders of magnitude faster than `FaST-LMM`. Even for 1 000 samples, 10^6^ SNPs and 500 permutations `permGWAS` (GPU) takes less than 1.8 h. In contrast, `EMMAX` would require at least more than 2.7 days, while `FaST-LMM` might take more than 11 days for 500 permutations. Results for environments with 8 cores are summarized in the Suppl. Fig. 1 and show similar results. Additionally, we over 500 *Arabidopsis thaliana* phenotypes with 100 permutations each on a single GPU (Nvidia RTX A5000 with 24GB RAM) in less than 8 days. The respective runtimes are shown in Supplementary Fig. 2. Notable, for phenotypes with a sample size above 800 individuals, the 24GB RAM weren’t sufficient and the analyses has been performed on an HPC environment allowing for additional RAM.

In summary, `permGWAS` is more efficient than the commonly used state-of-the-art LMMs, such as `EMMAX` and `FaST-LMM`, due to its tensor-based and batch-wise reformulation. Especially when performing GWAS with more than a few hundred of samples and permutations, `EMMAX` and `FaST-LMM` take several days to weeks to compute the results, while our implementation only needs a few hours. Although the GPU implementation of `permGWAS` is faster than the corresponding CPU implementation still outperforms existing methods.

#### 3.1.2 False Discovery Rate for Skewed Phenotypes

Our simulations show that the phenotype-wise FDR increases, if the respective phenotypes become more skewed. Using a static Bonferroni threshold, the phenotype-wise FDR increases from 30% for slightly skewed phenotypes to 50% in the most extreme case (Fig. 3D). The latter means that in nearly all of the simulated phenotypes, not only true, but also false associations have been found (Suppl. Tab. 2).

**Figure 3:**
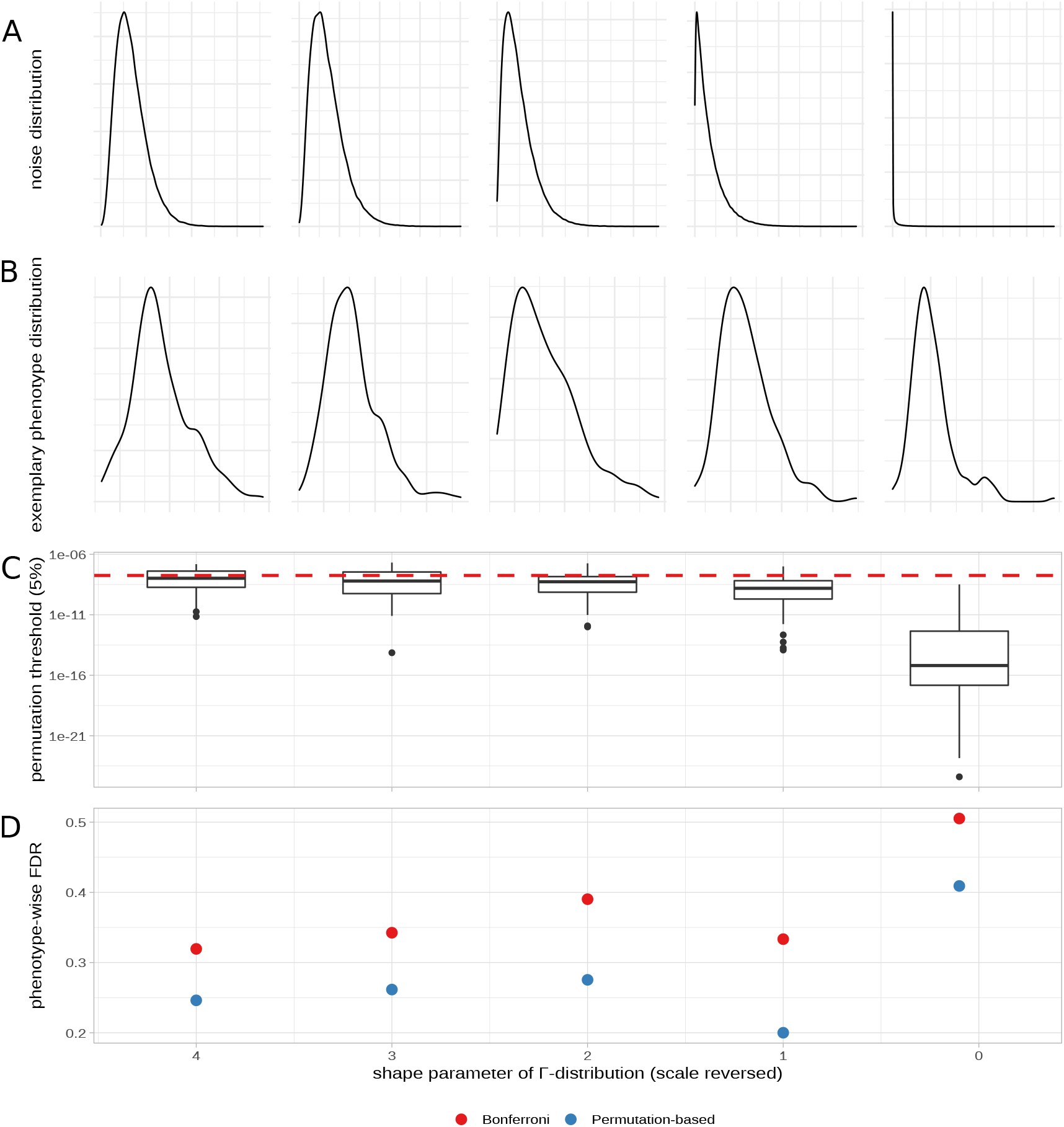
Simulated phenotypes with gamma-distributed noise. Shape parameters of the gamma distributions were set at 4, 3, 2, 1 and 0,1. (A) Shape of the gamma distribution. (B) Exemplary phenotypic value distribution for each shape parameter. (C) Permutation-based thresholds over 50 simulated phenotypes as box plots for each gamma shape parameter. Red dashed line illustrates the fixed Bonferroni significance threshold. (D) Phenotype-wise false discovery rate (FDR) for both the fixed Bonferroni significance threshold and the permutation-based significance threshold.

Noticeably, the permutation-based threshold becomes more and more stringent, if the phenotypic distribution becomes more skewed (Fig. 3C), thereby controlling the phenotype-based FDR more reliable (Fig. 3D and Suppl. Tab. 2). Skewed phenotypic distributions will violate model assumptions, and associations with low p-values can arise randomly. This will get controlled by permutations that can account for model violations, as underlying assumptions are also violated in a model without genetic signal. Hence, permutation can control for false associations that arise through non-normal phenotypic distributions.

On the other hand, for normally distributed phenotypes, the permutation-based threshold is less stringent compared to the Bonferroni threshold (Suppl. Fig. 3C) and increases the power to recover true associations (Suppl. Fig. 3D). However, also slightly more false positives are detected. To summarize, simulations suggested that a permutation-based threshold is more flexible, compared to a static Bonferroni threshold and will provide a higher power to detect true associations for normally distributed phenotypes, as well as control FDR for skewed phenotypes.

### 3.2 Permutation-based GWAS in Arabidopsis thaliana

After we highlighted the advantages of a permutation-based threshold with simulated data, we re-analyzed 516 real phenotypes that we have downloaded from the phenotypic data repository AraPheno (Seren *et al*., 2016; Togninalli *et al*., 2020). As expected for real data, many of these are non-normally distributed. Using the Shapiro-Wilk test on the phenotypic data, only 90 phenotypes had a p-value > 0.05, indicating a normal distribution (Suppl. File 1). As expected, by our simulations, we observed a correlation between the phenotypic distribution and the calculated permutation-based threshold (Suppl. Fig. 4). All, but two phenotypes that are normally distributed (Shapiro-Wilk test > 0.05), show a less stringent permuation-based threshold compared to the Bonferroni threshold (Suppl. Fig. 4 inset). In summary, for the 516 analyzed phenotypes, the permutation-based thresholds are 293 times more stringent and 223 less stringent compared to the Bonferroni threshold. Although, we don’t know the ground truth of true and false associations for this data, permutation-based thresholds markedly reduce the overall number of association, especially for skewed phenotypes. Comparing the 100 most skewed phenotypes (p-value from the Shapiro-Wilk test < 10^19^), nearly all (96) show a significant association using the Bonferroni threshold, while only six of the most normal distributed phenotypes (p-value from the Shapiro-Wilk test > 0.02) have a significant association. Using the permutation-based threshold these numbers change to 53 and 15, respectively (summary results of all analyses can be found in Suppl. File 1). *A priori*, there is no reason, why skewed phenotypes should more often show true association, therefore the number of reported associations with the permutation-based threshold seem more realistic. In general, we can observe different scenarios: (1) For some cases, a less stringent permutation-based threshold will identify a significant signal that would not have been significant using the Bonferroni threshold (Fig. 4A). This scenario is true for 22 different phenotypes, especially if their phenotypic distribution is normal (Suppl. Fig. 5A); (2) We observed 123 cases, where the Bonferroni threshold would indicate significant associations, but the permutation-based threshold would rather assume that these are false positives (Fig. 4B) and (3) for another 111 cases, even after using a permutation-based threshold, skewed phenotypes show still significant associations (Fig. 4C). Most phenotypes that belong to scenario (2) or (3) are non-normally distributed (Suppl. Fig. 5B and C).

**Figure 4:**
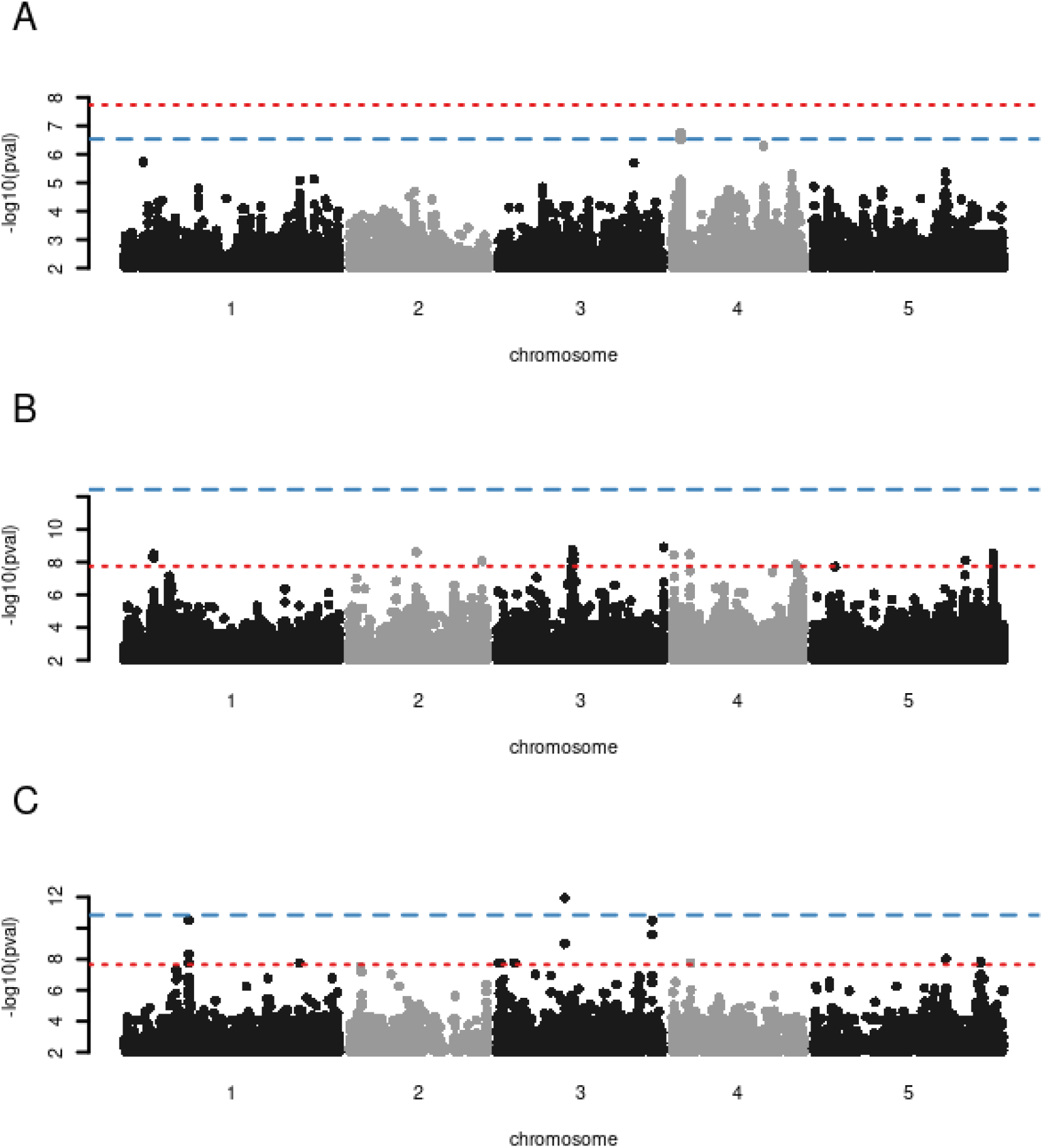
GWAS of three different *Arabidopsis thalaiana* phenotypes. Manhattan plots display the associations of all markers for the three phenotypes (A) #744 ((https://arapheno.1001genomes.org/phenotype/744/), which is nearly normal distributed, (B) #118 ((https://arapheno.1001genomes.org/phenotype/118/) and (C) #325 ((https://arapheno.1001genomes.org/phenotype/325/). Where the two latter phenotypes are non-normally distributed. The Bonferroni threshold is denoted by a red horizontal dashed line and the respective permutation-based threshold by a horizontal blue line.

Although, follow-up experiments would be needed to confirm that association deemed positive in the scenarios (1) and (3) are true positives, anecdotally many of this candidates seem plausible.

#### 3.2.1 Number of permutations and minor allele frequencies

In the previous paragraph we emphasized the benefit of using a permutation-based threshold, instead of a Bonferroni threshold. We performed 100 permutations for each phenotype, but more permutation could give a more accurate estimate of the threshold. To investigate the effect of the number of permutations on the permutation-based threshold, we performed additional permutations for two different Arabidopsis thaliana phenotypes. One phenotype is nearly normally distributed, while the other is markedly skewed (Suppl. Fig. 6A and C). We performed 100, 200, 300, 400, 500 and 1000 permutations. Here the used 5% threshold is nearly identical and stable across the different number of permutations performed for both phenotypes (Suppl. Fig. 6B and D). Thus, our empirical results suggest that 100 will give rise to a reliable estimate of the threshold and enable a fast analysis of many phenotypes and or huge data.

Next, we analyzed if minor allele frequency has an effect on false positives and the respective calculated permutation-based thresholds. It has been suggested that rare variants can easily associate with phenotypic extremes and thus that false positive associations of rare alleles are more prone in non-normally distributed phenotypes (Peloso *et al*., 2016). If this is true, a permutation-based threshold should be able to account for excessive false associations of rare alleles. Using `permGWAS` with increasing minor allele filters and thereby excluding rare alleles from the analysis, the Bonferroni threshold is just reflecting the lower amount of markers tested, while the permutation-based threshold has a non-linear dependency. For normally distributed phenotypes, the change in the permutation-based threshold is similar to Bonferroni (Suppl. Fig. 7A), while for a skewed phenotype a clear effect of excluding rare alleles is observed. As an example, in the analysis of phenotype #372 (DOI:10.21958/phenotype:372) from Arabidopsis thaliana, the calculated permutation-based threshold increases from 10^-16^ if all markers are analysed to 10^-10^ if only alleles with a minor allele count of at least 10 are considered (Suppl. Fig. 7B). For skewed phenotypes, the permutation-based threshold is clearly dependent on the allele frequency. `permGWAS` can compute and provide a distinct threshold for different allele frequencies that is - unlike Bonferroni - dependent on the phenotpic distribution and not the amount of markers tested.

## 4 Conclusions

We introduced `permGWAS`, an efficient linear mixed model for genome-wide association studies with population structure correction and permutation-based significance thresholds that can reliable control false positives for phenotypes with skewed distributions. Our method uses a 4D tensor reformulation of a linear mixed model using a permutation strategy proposed by Westfall-Young (Westfall and Young, 1993) to compute univariate association tests batch-wise, on both modern multi-core and GPU environments. We compared `permGWAS` in terms of runtime with `EMMAX` (Kang *et al*., 2010) and `FaST-LMM` (Lippert *et al*., 2011), two state-of-the-art linear mixed models. We could show that `permGWAS` outperformed both models in terms of computational and statistical performance. Especially, `permGWAS` is highly efficient in a permutation-based setting, due to the 4D tensor reformulation and the available GPU support (2h for `permGWAS` (`GPU`) vs. several days for `EMMAX` and `FaST-LMM` for 1 000 samples, 10^6^ markers and 500 permutations). These reformulations enable to perform permutation-based thresholds in practice.

We demonstrated through simulations and the re-analyses of public available data from the model plant species *Arabidopsis thaliana* that the use of a permutation-based threshold has many advantages compared to the classically used Bonferroni threshold. Bonferroni correction is thought as a very conservative way to control false positives in GWAS, and indeed for normal distributed phenotypes, we could show that the permutation-based threshold is less stringent and can identify more true positive associations. On the other hand, for non-normally distributed phenotypes, as often observed in biological data, the permutation-based threshold is quite often even more stringent. Here, our data suggest that we could reliably control false positives under those scenarios. To summarise, we highlight that the use of a permutation-based threshold should be considered the default choice in any GWAS and provide with `permGWAS` the tool to enable this.

## Author Contributions

AK and DGG conceived the study and supervised the project. MJ developed the mathematical framework. MJ implemented the model with input from JAF. MJA and CA simulated the phenotypes. MJ and MJA conducted the experiments. MJ, MJA, AK and DGG analyzed the results. MJ, AK and DGG wrote the paper with the contribution of all authors.

## 5 Supplementary Information

### Supplementary Tables

**Supplementary Table 1:** Mathematical Symbols and Notations

|  |  |
| --- | --- |
| $n$ | number of samples |
| $m$ | number of markers |
| $c$ | number of fixed effects |
| $b$ | number of batches |
| $q$ | number of permutations |
| $\sigma_g^2$ | extend of the genetic variance |
| $\sigma_e^2$ | extend of the residual variance |
| $\alpha$ | significance threshold |
| $\delta$ | corrected significance threshold for Family-wise error rate |
| $\delta^*$ | optimal significance threshold for FWER |
| $\delta_b^* = \frac{\alpha}{m}$ | Bonferroni adjusted significance threshold |
| $T_j$ | random variable for observed test statistic of the $j^{\text{th}}$ marker |
| $^{(k)}t_j$ | permutation test statistic of $j^{\text{th}}$ marker and $k^{\text{th}}$ permutation |
| $p_j$ | permutation based p-value of the $j^{\text{th}}$ marker |
| $\mathbb{1}$ | indicator function, taking the value 1 if the argument is true and 0 otherwise |
| $^{(k)}t_{\max} = \max_{j \in \{1, \dots, m\}} ^{(k)}t_j$ | maximal test statistic of $k^{\text{th}}$ permutation |
| $\tilde{p}_j$ | adjusted p-value for Westfall-Young permutation testing |
| $^{(k)}p_{\min}$ | minimal p-value of $k^{\text{th}}$ permutation |
| $RSS_0$ | residual sum of squares of the null model |
| $\mathbf{y} \in \mathbb{R}^n$ | vector of observed phenotypic values |
| $\mathbf{u} \in \mathbb{R}^n$ | vector of random effects |
| $\mathbf{x}_j \in \mathbb{R}^n$ | vector containing the $j^{\text{th}}$ SNP |
| $\boldsymbol{\epsilon} \in \mathbb{R}^n$ | vector of residual effects |
| $\boldsymbol{\beta} \in \mathbb{R}^c$ | vector of fixed effects coefficients |
| $\mathbf{X} \in \mathbb{R}^{n \times c}$ | matrix of fixed effects |
| $\mathbf{K} \in \mathbb{R}^{n \times n}$ | genetic relationship matrix (kinship) |
| $\mathbf{I} \in \mathbb{R}^{n \times n}$ | identity matrix of dimension $n$ |
| $\mathbf{V} \in \mathbb{R}^{n \times n}$ | variance-covariance matrix $\sigma_g^2 \mathbf{K} + \sigma_e^2 \mathbf{I}$ |
| $\mathbf{C} \in \mathbb{R}^{n \times n}$ | lower triangle matrix with $\mathbf{C}\mathbf{C}^T = \mathbf{V}$ |
| $\mathbf{X}_j \in \mathbb{R}^{n \times c}$ | matrix of fixed effects, containing a column of ones, the covariates and the $j^{\text{th}}$ SNP |
| $\mathbf{X}_j^b \in \mathbb{R}^{b \times n \times c}$ | 3D tensor containing $\mathbf{X}_j, \dots, \mathbf{X}_{j+b-1}$ |
| $\mathbf{Y}^b \in \mathbb{R}^{b \times n \times 1}$ | 3D tensor containing $b$ copies of $\mathbf{y}$ |
| $\mathbf{C}^b \in \mathbb{R}^{b \times n \times n}$ | 3D tensor containing $b$ copies of $\mathbf{C}$ |
| $\tilde{\mathbf{X}}_j^b \in \mathbb{R}^{b \times n \times c}$ | 3D tensor containing the transformed data $(\mathbf{C}^b)^{-1} \mathbf{X}_j^b$ |
| $\tilde{\mathbf{Y}}^b \in \mathbb{R}^{b \times n \times 1}$ | 3D tensor containing the transformed data $(\mathbf{C}^b)^{-1} \mathbf{Y}^b$ |
| $(\tilde{\mathbf{X}}_j^b)^T \in \mathbb{R}^{b \times c \times n}$ | 3D tensor containing the transposed matrices $(\mathbf{C}^{-1} \mathbf{X}_j)^T, \dots, (\mathbf{C}^{-1} \mathbf{X}_{j+b-1})^T$ |
| $\boldsymbol{\beta}_j^b \in \mathbb{R}^{b \times c}$ | coefficients of fixed effects for SNPs $j, \dots, j+b-1$ |
| $RSS_j^b \in \mathbb{R}^b$ | residual sums of squares for SNPs $j, \dots, j+b-1$ |
| $RSS_0^b \in \mathbb{R}^b$ | vector containing $b$ copies of the residual sums of squares of the null model |
| $t_j^b \in \mathbb{R}^b$ | test statistics of SNPs $j, \dots, j+b-1$ |
| $^{(k)}\sigma_g^2$ | genetic variance component of $k^{\text{th}}$ permutation |
| $^{(k)}\sigma_e^2$ | residual variance component of $k^{\text{th}}$ permutation |
| $^{(k)}\mathbf{y} \in \mathbb{R}^n$ | $k^{\text{th}}$ permutation of phenotype vector $\mathbf{y}$ |
| $^{(k)}\mathbf{V} \in \mathbb{R}^{n \times n}$ | variance-covariance matrix $^{(k)}\sigma_g^2 \mathbf{K} + ^{(k)}\sigma_e^2 \mathbf{I}$ of $k^{\text{th}}$ permutation |
| $^{(k)}\mathbf{C} \in \mathbb{R}^{n \times n}$ | lower triangle matrix with $^{(k)}\mathbf{C}^{(k)}\mathbf{C}^T = ^{(k)}\mathbf{V}$ |
| $^{(k)}\mathbf{Y}^b \in \mathbb{R}^{b \times n \times 1}$ | 3D tensor containing $b$ copies of the permutation $^{(k)}\mathbf{y}$ |
| $^{(k)}\mathbf{C}^b \in \mathbb{R}^{b \times n \times n}$ | 3D tensor containing $b$ copies of $^{(k)}\mathbf{C}$ |
| $^q\mathbf{X}_j^b \in \mathbb{R}^{q \times b \times n \times c}$ | 4D tensor containing $q$ copies of $\mathbf{X}_j^b$ |
| $^q\mathbf{Y}^b \in \mathbb{R}^{q \times b \times n \times 1}$ | 4D tensor containing $^{(1)}\mathbf{Y}^b, \dots, ^{(q)}\mathbf{Y}^b$ |
| $^q\mathbf{C}^b \in \mathbb{R}^{q \times b \times n \times n}$ | 4D tensor containing $^{(1)}\mathbf{C}^b, \dots, ^{(q)}\mathbf{C}^b$ |
| $^q\widetilde{\mathbf{X}}_j^b \in \mathbb{R}^{q \times b \times n \times c}$ | 4D tensor containing the transformed data $(^q\mathbf{C}^b)^{-1} {}^q\mathbf{X}_j^b$ |
| $^q\widetilde{\mathbf{Y}}^b \in \mathbb{R}^{q \times b \times n \times 1}$ | 3D tensor containing the transformed data $(^q\mathbf{C}^b)^{-1} {}^q\mathbf{Y}^b$ |
| $(^q\mathbf{X}_j^b)^T \in \mathbb{R}^{q \times b \times c \times n}$ | 4D tensor containing the transposed 3D tensors $(\mathbf{X}_j^b)^T, \dots, (\mathbf{X}_{j+b-1}^b)^T$ |
| $^q\boldsymbol{\beta}_j^b \in \mathbb{R}^{q \times b \times c}$ | coefficients of fixed effects for SNPs $j, \dots, j+b-1$ and $q$ permutations |
| $^q\mathbf{RSS}_j^b \in \mathbb{R}^{q \times b}$ | residual sums of squares for SNPs $j, \dots, j+b-1$ and $q$ permutations |
| $^q\mathbf{RSS}_0^b \in \mathbb{R}^{q \times b}$ | matrix containing $b$ copies of the residual sums of squares of the null model for each permutation |
| $^q\mathbf{t}_j^b \in \mathbb{R}^{q \times b}$ | test statistics of SNPs $j, \dots, j+b-1$ for $q$ permutations |

**Supplementary Table 2:** Count of true positive (TP) and false positive(FP) phenotypes out of 50 simulated phenotypes, and phenotype-wise false discovery rate (FDR) per effect strength.

| threshold | effect | measure | normal | 4 | 3 | 2 | 1 | 0.1 |
| --- | --- | --- | --- | --- | --- | --- | --- | --- |
| bonf | 1.0 | TP | 4 | 7 | 6 | 4 | 8 | 10 |
| bonf | 1.0 | FP | 3 | 7 | 13 | 19 | 22 | 49 |
| bonf | 1.0 | FDR | 0.43 | 0.50 | 0.68 | 0.83 | 0.73 | 0.83 |
| bonf | 1.5 | TP | 40 | 39 | 42 | 40 | 42 | 39 |
| bonf | 1.5 | FP | 7 | 14 | 19 | 25 | 23 | 49 |
| bonf | 1.5 | FDR | 0.15 | 0.26 | 0.31 | 0.39 | 0.35 | 0.56 |
| bonf | 2.0 | TP | 50 | 49 | 48 | 50 | 50 | 48 |
| bonf | 2.0 | FP | 17 | 23 | 25 | 32 | 25 | 49 |
| bonf | 2.0 | FDR | 0.25 | 0.32 | 0.34 | 0.39 | 0.33 | 0.51 |
| perm | 1.0 | TP | 5 | 10 | 7 | 3 | 2 | 1 |
| perm | 1.0 | FP | 7 | 5 | 5 | 5 | 3 | 13 |
| perm | 1.0 | FDR | 0.58 | 0.33 | 0.42 | 0.62 | 0.60 | 0.93 |
| perm | 1.5 | TP | 46 | 39 | 39 | 31 | 30 | 4 |
| perm | 1.5 | FP | 15 | 15 | 11 | 13 | 6 | 14 |
| perm | 1.5 | FDR | 0.25 | 0.28 | 0.22 | 0.29 | 0.17 | 0.78 |
| perm | 2.0 | TP | 50 | 49 | 48 | 50 | 48 | 26 |
| perm | 2.0 | FP | 20 | 16 | 17 | 19 | 12 | 18 |
| perm | 2.0 | FDR | 0.29 | 0.25 | 0.26 | 0.28 | 0.20 | 0.41 |

### Supplementary Figures

**Supplementary Figure 1:**
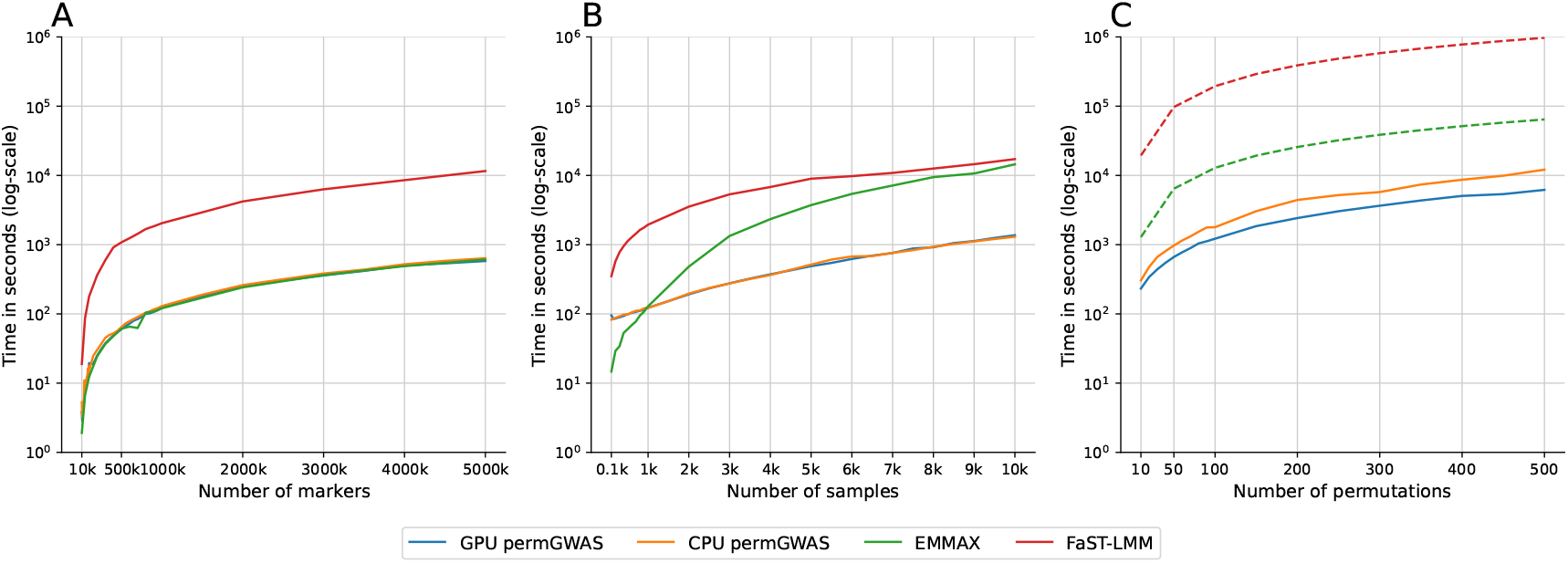
Runtime comparison of permGWAS vs. EMMAX and FaST-LMM using 8 cores. Note all axes are log-scaled. (A) Computational time as function of number of SNPs with fixed number of 1000 samples. (B) Computational time as function of number of samples with 10^6^ marker each. (C) Computational time as function of number of permutations with 1 000 samples and 10^6^ marker each. Dashed lines for `EMMAX` and `FaST-LMM` are estimated based on the computational time for 1000 samples and 10^6^ markers times the number of permutations.

**Supplementary Figure 2:**
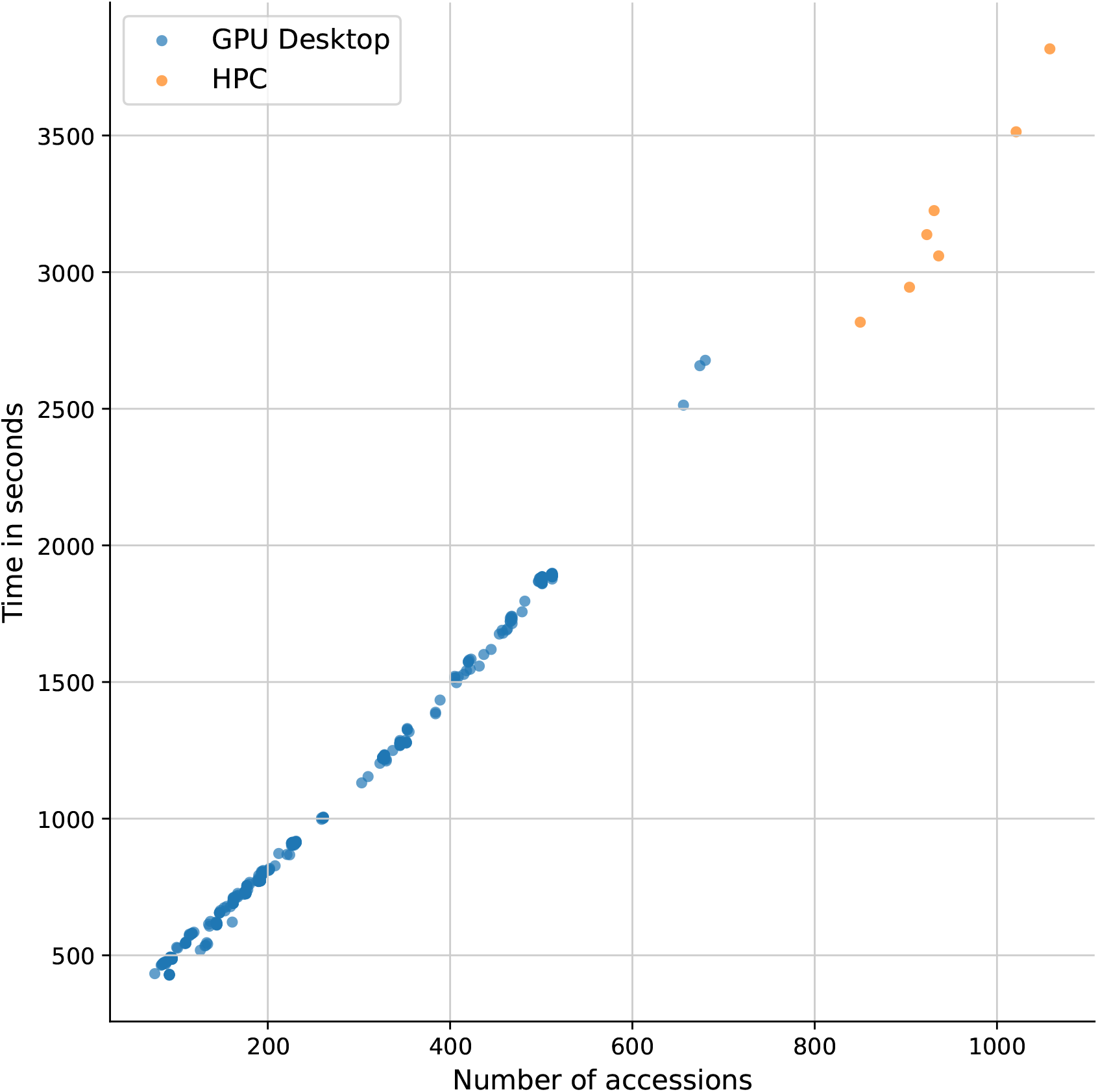
Runtime comparison of 516 phenotypes from *Arabidopsis thaliana* using 100 permutations each. Blue dots represent runtime of GWAS on a desktop machine with one Intel Xeon 8 core CPU with 3.5GHZ, 128GB of memory and a single NVIDIA RTX A5000 GPU with 24GB memory. Orange dots are runtime measurements of GWAS on a High Performance Cluster (HPC) including a NVIDIA A100 GPU.

**Supplementary Figure 3:**
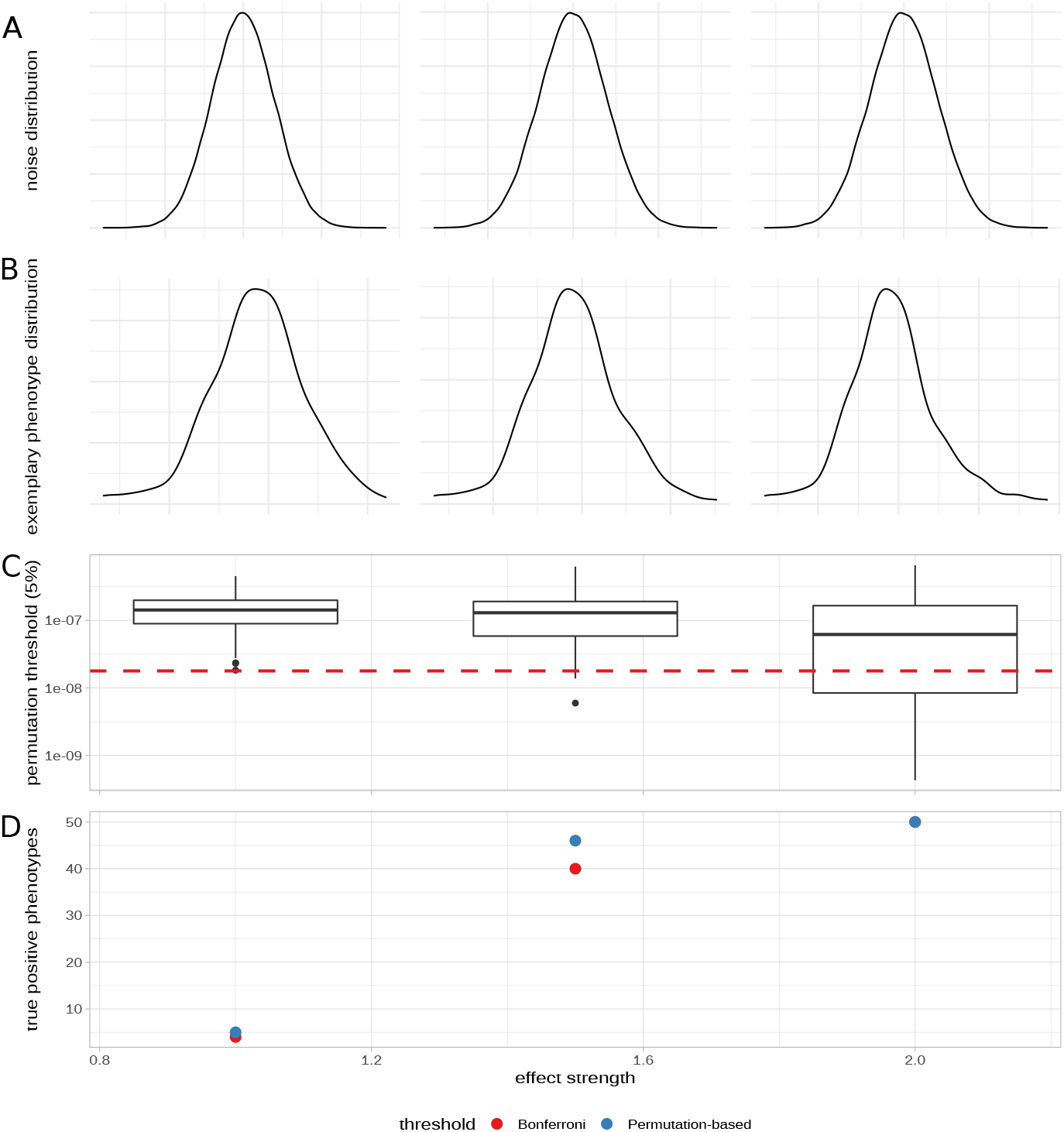
Simulated phenotypes with normally distributed noise and different effect strengths. In simulation the effect strength of the causative SNP was chosen to explain about 10% of total variance. The calculated phenotypic value was than multiplied by a factor (1.0, 1.5, or 2.0) to get different effect strengths. (A) Shape of the noise (normal). (B) Exemplary phenotypic value distribution for each shape parameter. (C) Permutation-based thresholds over 50 simulated phenotypes as box plots for each effect strength. Red dashed line illustrates the fixed Bonferroni significance threshold. (D) Number of true positive (TP) phenotypes (out of 50) for both the fixed Bonferroni significance threshold and the permutation-based significance threshold.

**Supplementary Figure 4:**
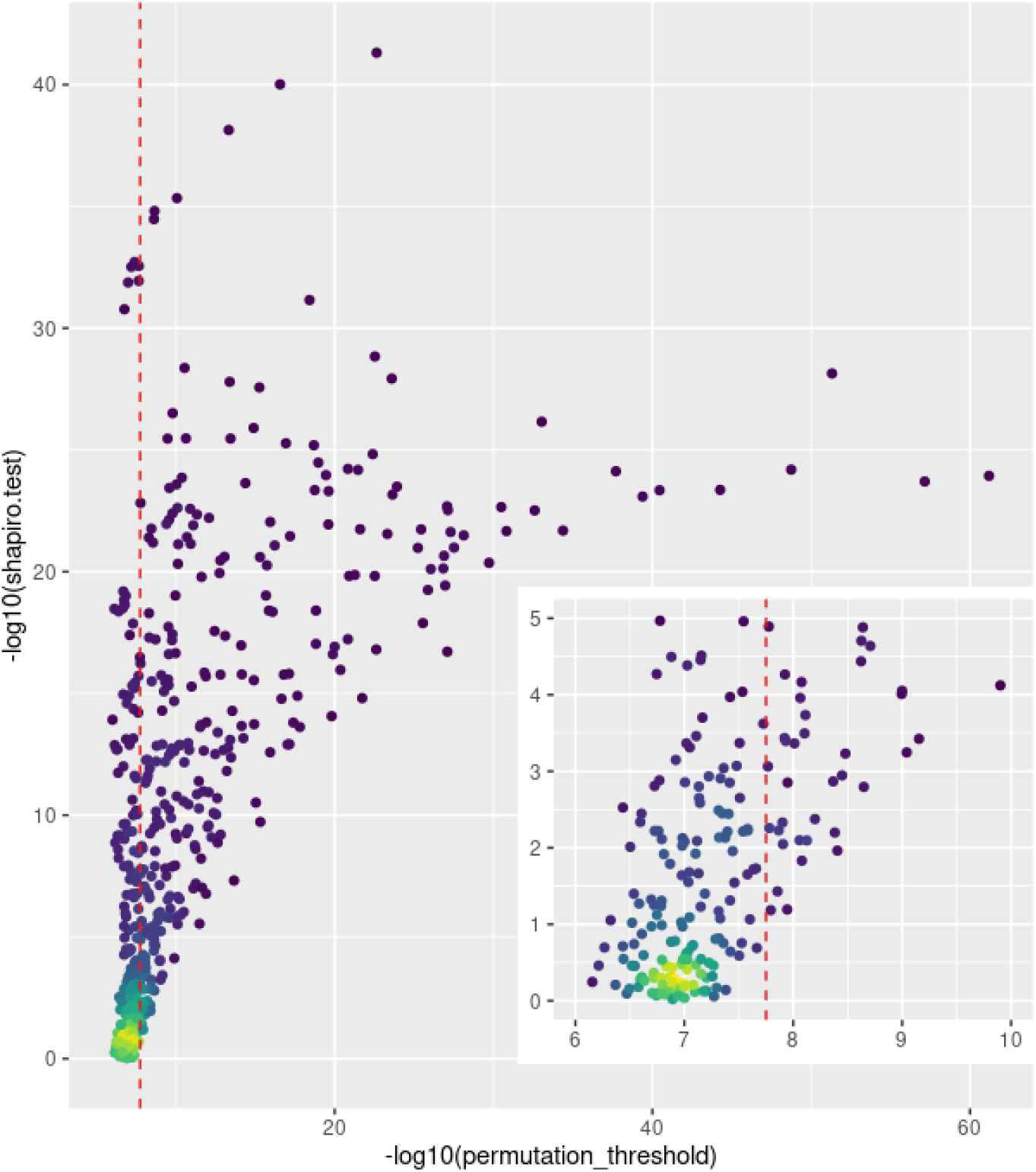
Correlation between permutation-based thresholds and the p-value from a Shapiro-Wilk test on the phenotypic distribution of 516 *Arabidopsis thaliana* phenotypes. The static Bonferroni threshold for 2.8 M markers is shown by a red vertical dashed line. Note that the shown threshold is calculated for 2.8 M markers and might differ slightly for phenotypes with small samples sizes. Each dot represents one phenotype and the false colors denote the amount of phenotypes at the same coordinates. The inset enlarges the region for normal and nearly normal distributed phenotypes.

**Supplementary Figure 5:**
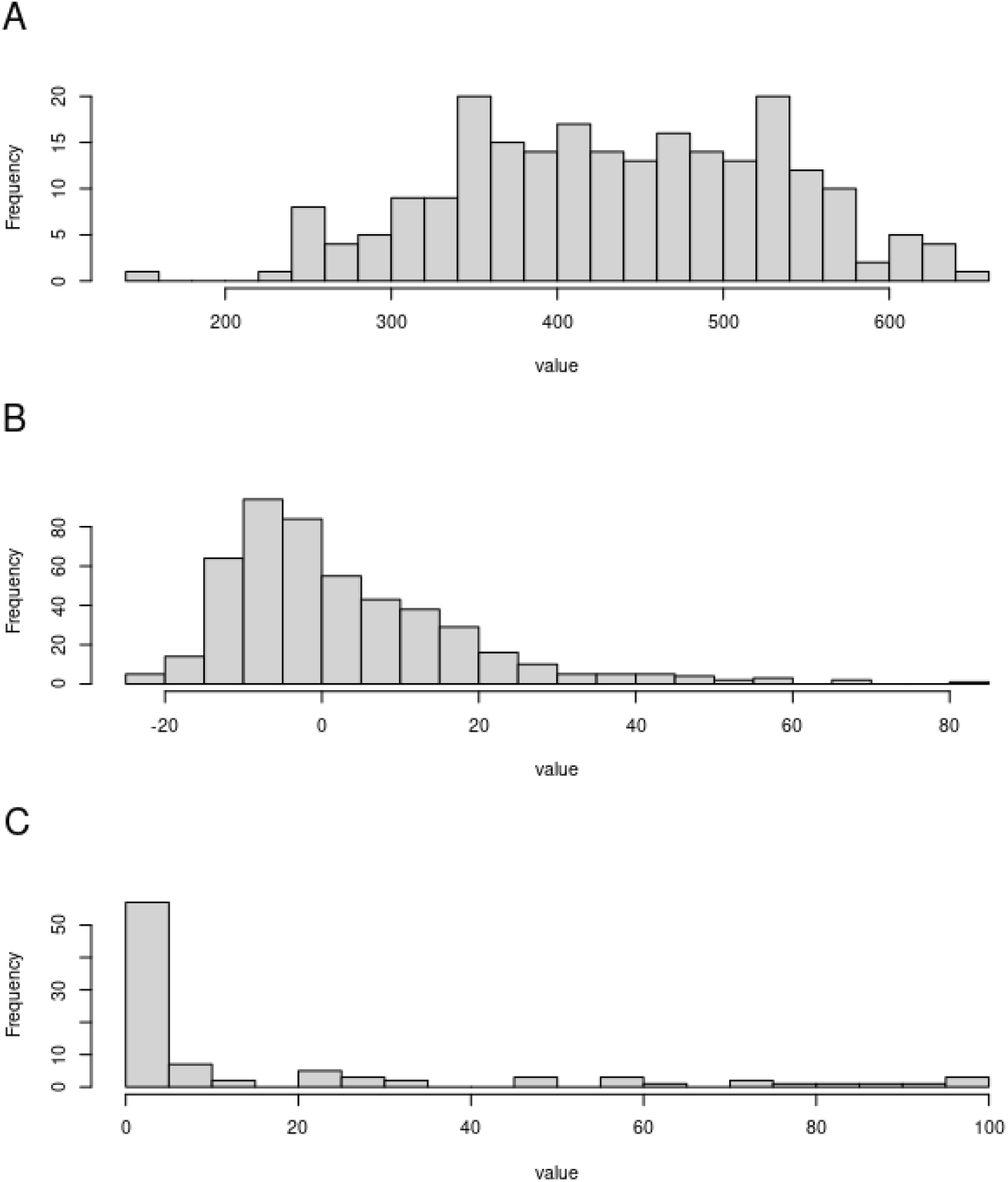
Histogram of the phenotypic distribution of three different *Arabidopsis thaliana* phenotypes. (A) Phenotype 744 (https://arapheno.1001genomes.org/phenotype/744/), which is nearly normal distributed (p=0.04)., (B) Phenotype 118 (https://arapheno.1001genomes.org/phenotype/118/), which is skewed and non-normally distributed (p<1e-17). (C) Phenotype 325 (https://arapheno.1001genomes.org/phenotype/325/), which is zero inflated and also non-normally distributed (p<1e-12).

**Supplementary Figure 6:**
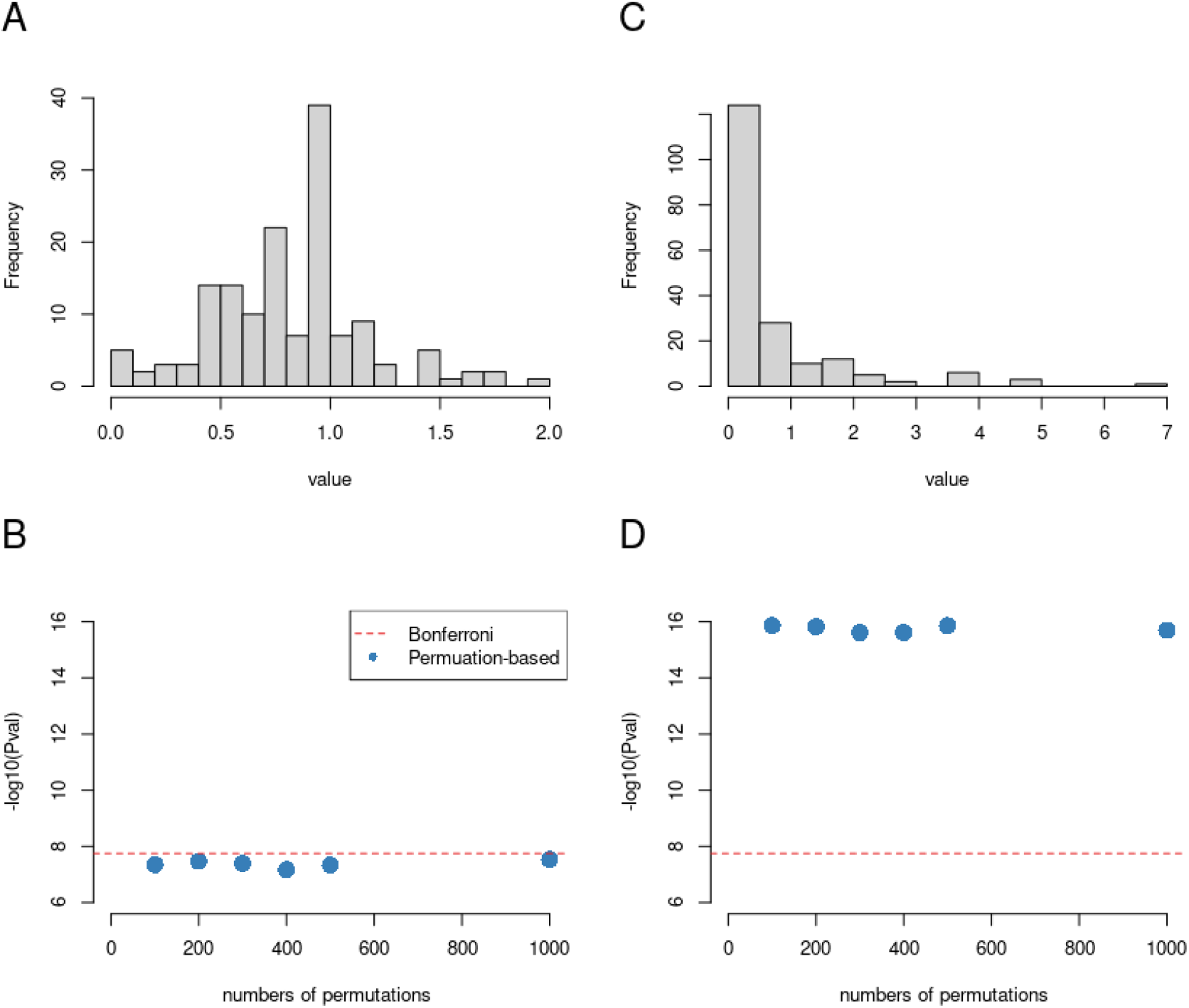
Permutation-based thresholds with different amounts of permutations. Two different *Arabidopsis thaliana* phenotypes have been analyzed with varying amounts of permutations. Phenotype 1271 ((https://arapheno.1001genomes.org/phenotype/1271/) is nearly normal distributed and shown in (A) and (B), while phenotype 372 ((https://arapheno.1001genomes.org/phenotype/372/) is non-normally distributed ((C) and (D)). The Bonferroni threshold is denoted by a red horizontal dashed line and the respective permutation-based thresholds for different numbers of permutations is shown by blue dots. Note that the latter is stable between 100 and 1000 permutations both for the normally and non-normally distributed phenotype.

**Supplementary Figure 7:**
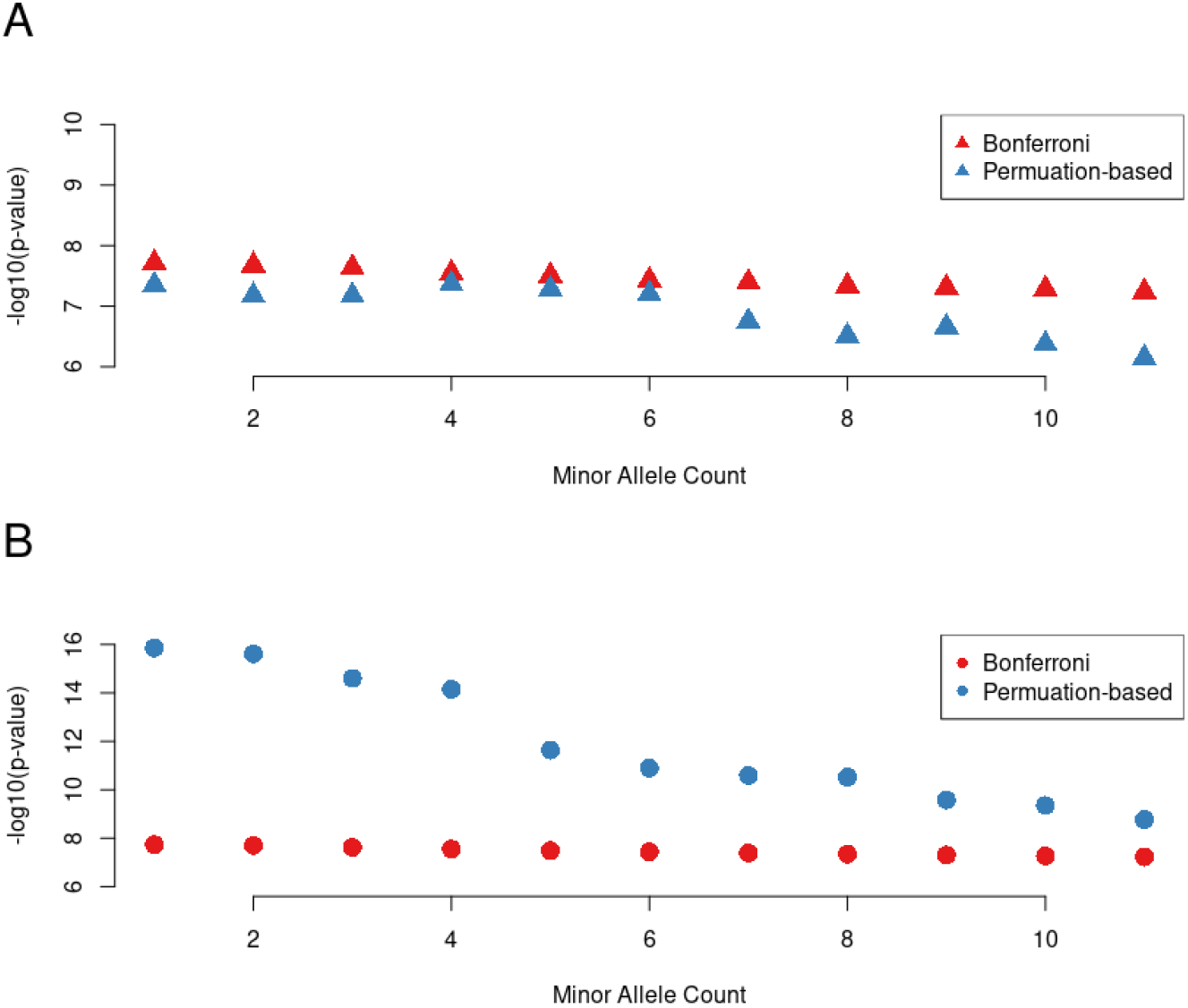
The effect of minor allele filters on Bonferroni and permutation-based thresholds. The respective thresholds are shown as a function of increasing minor allele filters, where rare alleles have been removed prior to the analysis. (A) shows a nearly normally distributed phenotype (1271) and (B) a non-normally distributed phenotype (372). The Bonferroni threshold (red) becomes slightly higher with an increasing minor allele filter, as fewer markers are tested. This increase is more pronounced for the permutation-based threshold, especially for the non-normally distributed phenotype. Here the increase in the threshold is non linear, but specific for the phenotypic distribution.

### Supplementary Files

**Supplementary File 1**: File includes permutation based thresholds and number of hits for all 516 AraPheno phenotypes. Supplementary files can be found at: https://github.com/grimmlab/permGWAS/tree/main/suppl_data.

